# Variability of immune gene expression among different groups within ant colonies show multifaceted response to infection by a non-lethal ectoparasitic fungus

**DOI:** 10.1101/2024.02.08.579503

**Authors:** Kincső Orbán-Bakk, Eva Schultner, Jürgen Heinze, Bálint Markó, Enikő Csata

## Abstract

Social insect colonies are known to be targeted by a wide variety of different parasites and pathogens because of their high host abundance. However, within a colony, the level of risk to exposure could vary among individuals depending on their role. Unlike many known parasites, which mostly target specific groups of individuals, e.g. foragers, the myrmecoparasitic fungus *Rickia wasmannii* infects entire ant colonies, being linked to subtle changes in physiology, morphology and behaviour. We investigated how different groups within the colonies respond to being exposed to the fungus by measuring the expression of the genes *defensin 1* and *prophenoloxidase*, both vital components of ant immunity. We found that workers, queens and broods varied in their immune response. Workers displayed diverse profiles, with variable responses to infection: in same-age workers, both *prophenoloxidase* and *defensin 1* levels exhibited increases in correlation with pathogen loads. Queens exhibited a more pronounced immune response. Highly infected queens had a heightened immune response. Larvae did not show a discernible response. Morphological and physiological characteristics had limited effects on gene expression, except in the case of queens, where larger individuals displayed higher *defensin 1* expression. Our study shows that these divergent responses likely stem from the differing physiological needs and priorities of various groups within the colony.

**Highlights:**

1. In same-age workers, *prophenoloxidase* and *defensin 1* levels increased with pathogen loads.
2. Body size affected *defensin 1* expression in a caste-specific manner: larger queens displayed higher expression.
3. Infection did not elicit any specific response in larvae.
4. The diverse response to infection likely arise from distinct physiological needs and priorities within colony groups.

## Introduction

Infection-activated immune genes play a crucial role in the defence of animals against pathogens. The immune response involves a complex interplay between innate and adaptive immunity (Cremer & Sixt 2009), each comprising a diverse array of genes to combat pathogens (Iriti & Faoro 2007). The innate immune response involves the production of antimicrobial molecules, like peptides, lysozymes, and complement proteins, which directly target and neutralise pathogens (Rosales 2017, Sharrock & Sun 2020). It also recruits immune cells to the infection site, aiding in pathogen containment and elimination (Akira et al. 2006, Dubovskiy et al. 2016). Additionally, infection-activated immune genes trigger the production of pro-inflammatory molecules, such as cytokines and chemokines, to coordinate the immune response and attract more immune cells to the infected area (Weber et al. 2003). In insects, the innate defence response involves different pathways, like Toll and Imd, which lead to the expression of antimicrobial peptides like defensin, and prophenoloxidase (Levashina 2004, Lemaitre & Hoffmann 2007, Orbán-Bakk et al. 2023).

Social insect colonies represent a rich resource for parasites and pathogens because of high abundance of host organisms, which are also genetically closely related allowing infections to spread both vertically from parent to offspring and horizontally within the same generation (Sherman et al. 1988, Schmid-Hempel 1998, Cremer et al. 2007). Ants try to hinder the spread of pathogens within the colony by various means, e.g., disposal of refuse material, exclusion of infected nestmates, changing the topology of the social network, and relocation of colonies to safer places (Howard & Tschinkel 1976, Schmid-Hempel 1998, Hart & Ratnieks 2001, Cremer et al. 2007, Konrad et al. 2012, Stroeymeyt et al. 2018). Parallel to these social measures, ants also respond individually by eg. activating their immune system to impede pathogen invasion on individual level (Schmid-Hempel & Ebert 2003; Konrad et al. 2015). Several studies have investigated the relationship between exposure to infection and immune gene expression in ants (Yek et al. 2013, Konrad et al. 2015, de Bekker et al. 2017, Stoldt et al. 2021). However, most of the studies have focused on workers, while ignoring the other two core components of colonies: queens and brood, specifically larvae. Thus, our understanding of a colony’s immune reaction to infection is incomplete.

The ectoparasitic fungus *Rickia wasmannii* Cavara, 1899 (Ascomycota, Laboulbeniales) can infect entire colonies of the host ant *Myrmica scabrinodis* Nylander 1846, being linked to noticeable changes in individual and social behaviour (Csata et al. 2014, Báthori et al. 2015, Csata et al. 2017a, Csata et al. 2023), as well as in morphology and physiology (Csata et al. 2018, Csősz et al. 2021, Orbán-Bakk et al. 2022). The infection is persistent and shortens the lifespan of infected individuals on the long run (Csata et al. 2014). Besides workers of all age classes, queens can also be heavily infected (Tartally et al. 2007, Figure 1). Larvae are not known to be infected though. However, since queens and young workers typically spend their time on or near the brood, this implies that in infected colonies larvae are also exposed to the threat of infection, which might trigger immunological changes.

**Figure 1.**
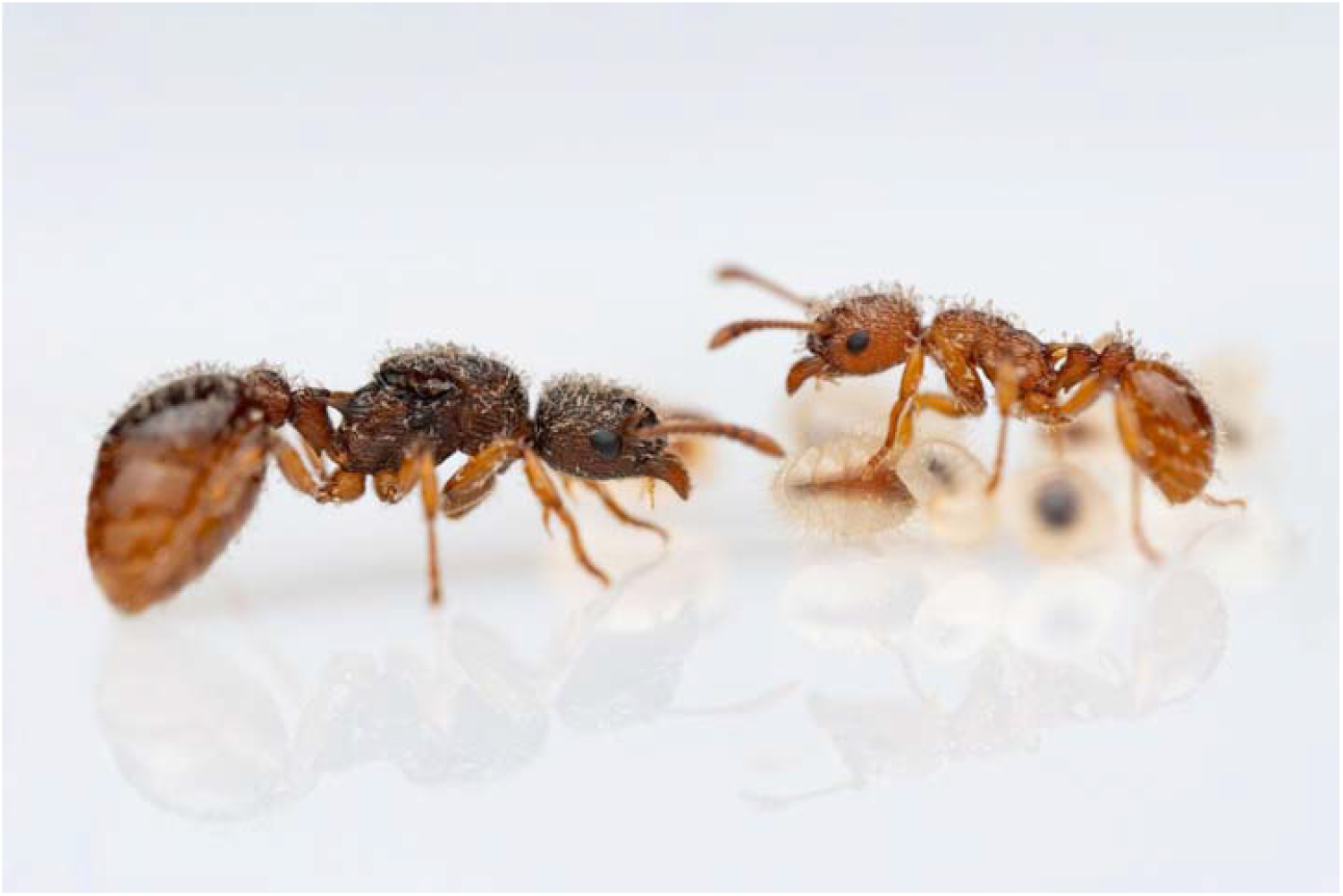
A heavily infected *Myrmica scabrinodis* queen with brood and an infected young worker. The whitish fur-like coverage on the body of the queen and worker are the thalli of *Rickia wasmannii* (photo by Daniel Sánchez García).

In the frame of this study, we assessed the immune response of queens, workers and larvae from infected and uninfected colonies. We measured the expression of the immune genes *defensin 1* and *prophenoloxidase*, both crucial components of humoral and cellular defences in ants (Konrad et al. 2015). *Prophenoloxidase* plays a vital role in the melanization cascade, crucial for encapsulating infectious agents in innate immunity. *Defensin* has an antibacterial and fungicidal effect (Bulet & Stöcklin 2005) and acts against Gram-positive bacteria (Viljakainen & Pamilo 2005). Considering the central role of the queen in a colony, and its considerably longer lifespan compared to workers, we hypothesized that their immune response would be more pronounced. Furthermore, we investigated whether exposure to the risk of infection can activate the immune defence of larvae. In adults, we also measured individual infection load, body size and fat content to reveal additional factors that might explain immune gene expression patterns. Overall, we predicted that even when faced with a non-lethal pathogen, ants upregulate immune gene expression, indicating a robust innate immune response to fungal parasites.

## Material and methods

### Field sampling and experimental design

Colonies of *M. scabrinodis* were collected near Bor□a Catun (N 46.88909, E 23.70067), Central Romania, from a North-exposed slope of a meso-xeric grassland (Czekes et al. 2012), where infected and uninfected colonies co-occur. The collected colonies belong to the same haplotype (Csata et al. 2023), thus no interference is expected from potential cryptic speciation in *M. scabrinodis* (Ebsen et al. 2019). In total, eight infected and seven uninfected colonies were collected at the end of July 2019. Fungal infection and the species of the ant was primarily identified with a hand magnifying glass in field conditions and later confirmed in laboratory conditions using an Olympus SZ51 stereomicroscope at ×80 magnification. *Myrmica scabrinodis* was identified based on the identification key of Czekes et al. (2012).

*Rickia wasmannii* Cavara (1899) stands out as the most prevalent ectoparasitic myrmecophilous species within the Holarctic region obligatorily exploiting *Myrmica* ants (Tartally et al. 2007, Lapeva-Gjonova and Santamaria 2011, Santamaria & Espadaler 2012, Csata et al. 2013, Witek et al. 2014, Haelewaters et al. 2015). The thalli of *Rickia wasmannii* attach to the outer layer of the cuticle, manifesting on the hosts’ surface in the form of club-shaped, setae-like structures (Haelewaters 2012, Tragust et al. 2016). Fungal thalli of *R. wasmannii* are easy to recognize as infected individuals appear unusually hairy (Figure 1).

In total, 14 infected and 12 uninfected experimental colonies were established from the field colonies (Table S1). Each of the experimental colonies contained one queen, 70 old workers (identified based on their dark brown cuticle; *see* Moron et al. 2008, Csata et al. 2017b), 15 to 30 worker pupae and 15 to 30 larvae. All experimental colonies were kept in artificial nests (15cm × 10cm × 5cm) under constant laboratory conditions (min. temperature = 12°C, max. temp. = 18°C, 12:12 light:dark cycle), and fed with an artificial carbohydrate- and protein-rich diet (*see* Bhatkar & Whitcomb 1970) three times per week.

The survival of the ants was monitored daily. To maintain experimental colony size, dead workers and hatched pupae were replaced from respective stock colonies. To avoid any influence of age on immune gene activation in workers, two days after hatching, young workers were marked with an individual combination of colour dots (©UNI Paint Markers) on the thorax and then followed each for a standard period (103 days) until they reach full maturity (dark brown cuticle). At the end of the standard period, they were collected and stored for analyses (Table S1). Altogether, 163 freshly hatched workers were marked (54 uninfected, 109 infected), out of which 43 survived (17 uninfected, 26 infected) during the standard observation period. At the time of worker sampling (103 days), 15 queens (7 uninfected, 8 infected), and 17 groups of three larvae (8 from uninfected colonies, 9 from infected colonies) were also collected and stored for later analyses (*see* Table S1).

### Gene expression assays

Prior to any analyses, the sting and venom gland were removed from each marked worker and queen as venom can interfere with qPCR (Bouzid et al. 2014). The gaster, thorax, and head of each individual were then separated and moved to individual tubes, flash-frozen in liquid nitrogen, and stored at -80°C. The larvae were pooled in groups of three, and also flash-frozen in liquid nitrogen, and stored at -80°C until RNA extraction. In insects, including ants, immune genes are expressed in the gut and the fat body (Hoffmann 1995; Gillespie et al. 1997; Hoffmann 2003; Shia et al. 2009; Tsakas & Marmaras 2010; Choppin et al. 2021). For this reason, only the gaster of workers and queens were used for gene expression assays, while the other body parts were used to assess the intensity of infection, body size (head), and fat content (thorax).

Immediately before RNA extraction, each sample tube was dipped into liquid nitrogen and then the ant was homogenized using a pestle. In the case of larvae, pooled samples of 3 individuals were used. RNA was extracted using the ReliaPrep RNA Cell Miniprep System (Promega, USA) according to the manufacturer’s protocol. RNA concentration was measured with a Qubit Fluorometer (Life Technologies, USA), and 100 ng of RNA were used for cDNA synthesis with the iScriptTM gDNA Clear cDNA Synthesis Kit (Bio-Rad Laboratories, USA), resulting in a final concentration of 5 ng/μl for each sample.

The expression levels of two insect immune genes were analyzed by real-time quantitative PCR (qPCR): *prophenoloxidase* (*PPO*), and *defensin 1* (*Def1*). Specific primers for *prophenoloxidase, defensin 1*, as well as a housekeeper (Y45) were designed BLASTing sequences obtained for *M. scabrinodis* (*Def1*), the European honey bee *Apis mellifera* (*PPO, AY242387.2*, www.hymenopteragenome.org), and the ant *Cardiocondyla obscurior* (*Y45*, Cobs_04843, Schrader et al. 2014, www.antgenomes.org) against the *M. scabrinodis* transcriptome (Global Ant Genomics Alliance (GAGA)). Intron-spanning primers were designed using Geneious Prime (version 2020.0.3) and tested on pools of larvae in a heat-gradient PCR. PCR products were purified and Sanger-sequenced (LGC Genomics, Berlin, Germany), and sequences were aligned in Geneious Prime to confirm primer specificity. Primer efficiencies were measured using a 5-step dilution series (Table 1). For each sample and gene, qPCR reactions were run in triplicates, from which we calculated mean Cq values. The expression of target genes relative to the housekeeper was assessed using the ΔCq method described by Livak and Schmittgen (2001). We obtained analysable quantities of RNA for 23 workers (10 infected, 13 uninfected), 15 queens (8 infected and 7 uninfected) and 17 pools of larvae (8 from uninfected colonies, 9 from infected colonies).

**Table 1.**
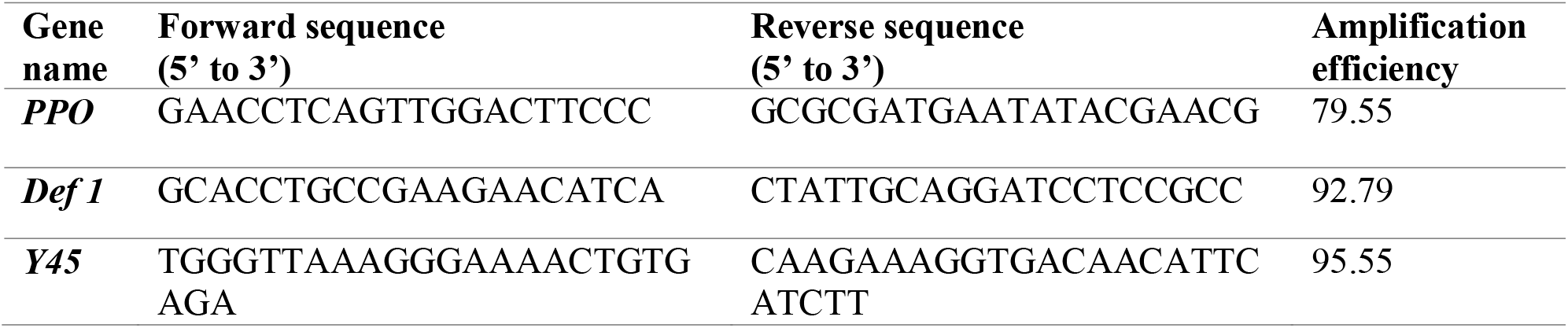
Primer sequences used for qPCR and amplification efficiencies (*PPO* = prophenoloxidase, *Def 1* = defensin, *Y45* = housekeeper gene).

For RNA extraction we used the ReliaPrep RNA Cell Miniprep System (Promega Kit). For the preparation of the Column Wash Solution, 9.5 ml of 95% ethanol was added to 5 ml of concentrated column wash solution. As a start of the protocol after grinding with a pestle, 250 μl BL+TG Buffers were added for each sample. We centrifuged (Eppendorf Microcentrifuge 5415R) the samples at room temperature for 1 min at 16000 x g and we removed 200 μl of supernatant and transferred it to a new 1.5 ml tube. After that, we admitted 200 μl isopropanol and vortexed for 5 sec. For the next steps, we followed the manufacturer’s instructions from the original Promega protocol. After applying the 30 μl of DNaseI master mix, we incubated the samples at room temperature for 45 min. Next, we added 200 μl of column wash solution and centrifuged at room temperature for 30 sec at 13000 x g. Furthermore, we added 500 μl RNA wash solution and centrifuged at room temperature for 30 sec at 13000 x g. After centrifugation, we added 300 μl RNA wash solution and centrifuged the samples again at room temperature for 2 min at 16000 x g. In the end, we added 30 μl Nuclease-free water to the sample, incubated it at 8 min, and then centrifuged it at 1 min at 13000 x g. As a last step, we incubated the sample at 1 min, and at least centrifuged at 13000 x g for 1 min.

### Infection intensity and body size

To quantify the intensity of infection, all fungal thalli were counted on the right side of each mature individual’s head with a Leica stereomicroscope at ×80 magnification. The head is the most infected body part of the ants in the case of *Rickia wasmannii* (Markó et al. 2016). An ocular micrometer was used to set the axial line through the ant’s head to separate the right and left sides. Larvae were not included in the analyses.

To detect relationships between body size and the level of immune gene expression (Castella et al. 2010), head sizes of queens and workers were also measured as an appropriate proxy for body size in ants (Porter 1983). A digital image of each head was captured and measured using a stereomicroscope (Keyence OP-87270) at ×150 magnification. Head size (HS) was calculated as head width (HW) × head length (HL) (μm^2^). Larvae were omitted from this analyses.

### Fat content analyses

The fat content of the thorax of each adult individual was measured to detect if there is any relationship between physiological conditions and immune gene expression. Each sample was dried at 60°C in an oven (Memmert U10) for five days and weighed to the nearest 0.0001 mg with a Sartorius SC2 ultra microbalance to measure its dry mass. The sample was then placed in 2 ml petroleum ether (boiling range 40-60°C, Merck, Darmstadt, Germany) at room temperature for two days, after which it was transferred to a new tube. The petroleum ether was renewed with 2 ml as in the previous step and the sample kept for two further days. The sample was then dried at 60°C for six days and weighed again to determine its lean mass. Fat content (%) was calculated according to the standard equation: (dry mass - lean mass) × 100/dry mass (e.g., Bernadou et al. 2015, Csata et al. 2017, Orbán-Bakk et al. 2022). As whole bodies of larvae were used for RNA extraction, we could not assess larval fat content.

## Statistical analyses

All data were tested for assumptions of normality. The response variables (2^-ΔCq^ values) that did not fit linear model requirements were transformed using the ‘*bestNormalize*’ function in the ‘*bestNormalize*’ package (Peterson and Cavanaugh 2019). To test for an overall correlation between expression levels (2^-ΔCq^ values) of *prophenoloxidase* and *defensin 1*, we performed a Spearman correlation using all data (function ‘*cor. test*’). Spearman correlations between the expression levels of the two genes were also calculated separately for workers, queens, and larvae.

To examine whether castes show different immune responses, we performed a linear mixed effect model (LMM). The model was run separately for each gene, using normalized 2^-ΔCq^ values as response variables and castes (larvae, workers and queens) as explanatory variables. Adjusted p-values were obtained using the ‘*emmeans*’ function in the ‘*emmeans*’ package (Russell 2023).

To investigate the effects of infection intensity, size, and fat content on immune gene expression in queens and workers, a linear mixed effect model (LMM) was run separately for each gene and caste using normalized 2^-ΔCq^ values as response variables and number of thalli, head size, and fat content and infection status of the individuals (infected or uninfected) as explanatory variables. For larvae, only the infection status of the colony was used as an explanatory variable, as they do not have thalli on their cuticle and because we did not assess larval fat content or size. In each model, the colony tag (the identity of the field colony) was entered as a random factor to handle dependencies. In each of the analyses, we obtained a minimal model by successively removing non-significant variables by stepwise backward elimination (threshold = 0.05).

All statistics were performed using R 4.1.0 (R Core Team, 2020). The models’ quality was assessed using residual diagnostics from the ‘*DHARMa*’ package (Hartig & Hartig 2017). This involved checking for deviations from the expected distribution, evaluating dispersion and outliers, and creating a plot of residuals against predicted values.

LMMs were applied using the ‘*lmer*’ function in the ‘*lme4*’ package (Bates et al. 2015). Test outputs (χ^2^ and p-values) were obtained using the function ‘*Anova*’ in the ‘*car*’ package (Fox and Weisberg 2020). The graphs were produced using the ‘*ggplot2*’ R package (Wickham 2009).

## Results

There was no correlation between the expression levels of *prophenoloxidase* and *defensin 1* when all data were considered together (Spearman correlation, rho = 0.05, p = 0.74), nor when each group was analysed separately (Spearman correlation, larvae: rho = 0.25, p = 0.33; workers: rho = 0.14, p = 0.61; queens: rho = 0.45, p = 0.16).

Overall, *prophenoloxidase* expression varied relatively little between castes (LMM, larvae vs queens: t = -0.1, p = 0.99; larvae vs workers: t = -1.73, p = 0.2; queens vs workers: t = -1.47, p = 0.31, Figure 2). Colony infection status did not affect *prophenoloxidase* expression in larvae (LMM, χ^2^ = 0.05, p = 0.94). Neither did in workers (LMM, colony infection status, χ^2^ = 1.6, p = 0.2), similarly to fat content (χ^2^ = 2.4, p = 0.12), and head size (χ^2^ = 2.5, p = 0.11). However, workers carrying more thalli showed higher *prophenoloxidase* expression (χ^2^ = 4.18, p = 0.04). In queens, almost all included variables had a significant effect on *prophenoloxidase* expression, except head size (LMM, χ^2^ = 0.13, p = 0.71). The number of thalli (χ^2^ = 7.8, p = 0.005), fat content (χ^2^ = 5.75, p = 0.01) and colony infection status (χ^2^ = 6.67, p = 0.009) were associated with increased *prophenoloxidase* expression (Figure 2).

**Figure 2.**
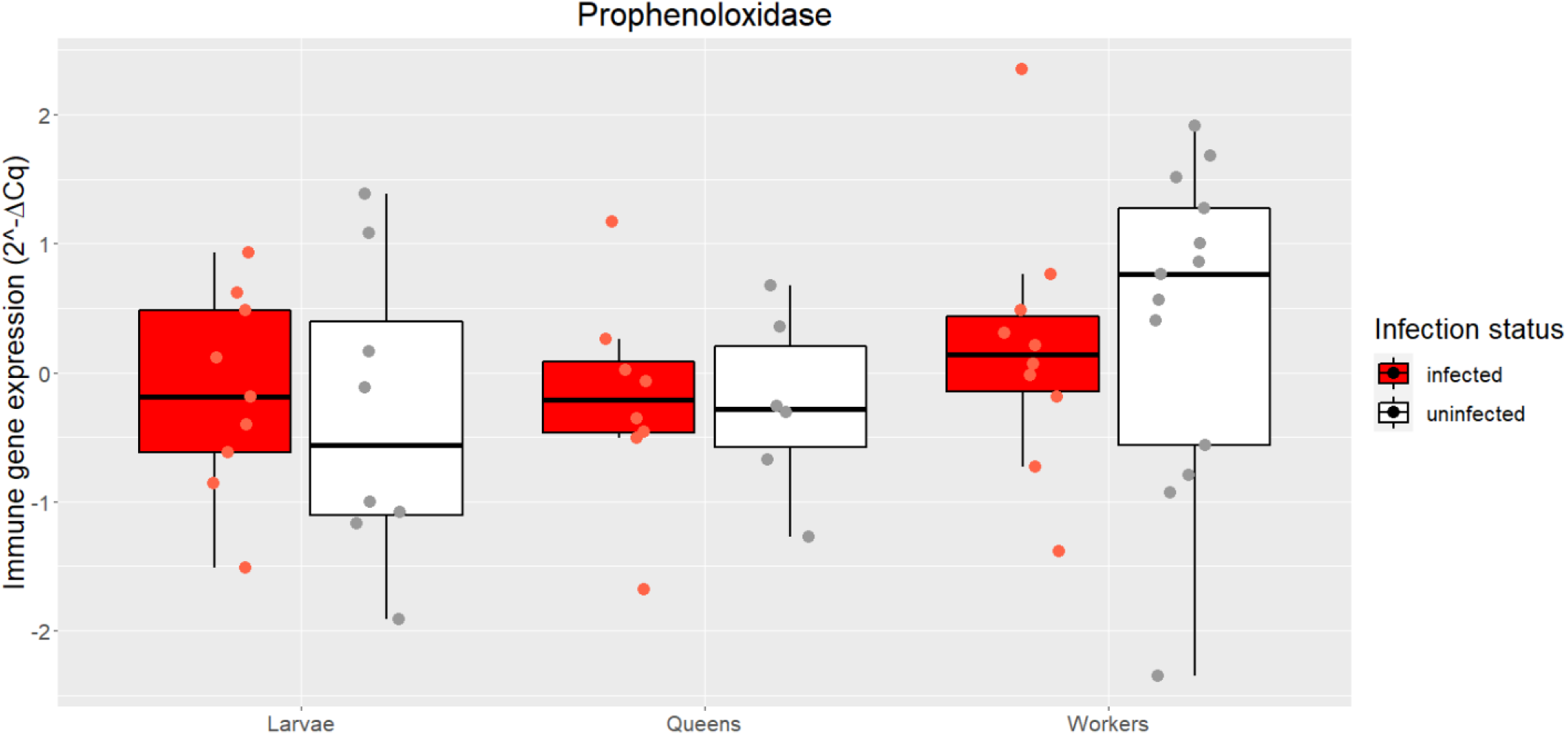
*Prophenoloxidase* expression levels (2^-ΔCq^ values) in larvae, queens, and workers from infected (red) and uninfected (white) colonies (transformed data according to the applied statistical analyses). Box-and-whisker plots are shown with median, 25% –75% quartiles.

In contrast to *prophenoloxidase, defensin 1* expression was highly variable across groups, with larvae showing higher *defensin 1* expression compared to queens and workers (LMM, larvae vs queens: t = 8.03, p < 0.001; larvae vs workers: t = 6.24, p = < 0.001; queens vs workers: t = - 2.11, p = 0.10, Figure 3). *Defensin 1* expression in larvae was not affected by colony infection status (LMM, χ^2^ = 0.09, p = 0.75). Neither did in workers (LMM, χ^2^ = 2.01, p = 0.15), similarly to head size (LMM, χ^2^ = 0.07, p = 0.78), and fat content (LMM, χ^2^ = 1.53, p = 0.21). However, workers carrying more thalli showed higher *defensin 1* expression (LMM, χ^2^ = 7.67, p = 0.005). The situation was quite different in queens. Thus, queens from infected colonies exhibited higher *defensin 1* expression (LMM, χ^2^ = 3.91, p = 0.04, Figure 3), whereas large queens expressed more *defensin 1* (LMM, χ^2^ = 27.61, p < 0.001). Neither the number of thalli (LMM, χ^2^ = 0.37, p = 0.54) nor fat content (LMM, χ^2^ = 3.08, p = 0.07) affected *defensin 1* expression in queens (LMM, χ^2^ = 3.08, p = 0.07).

**Figure 3.**
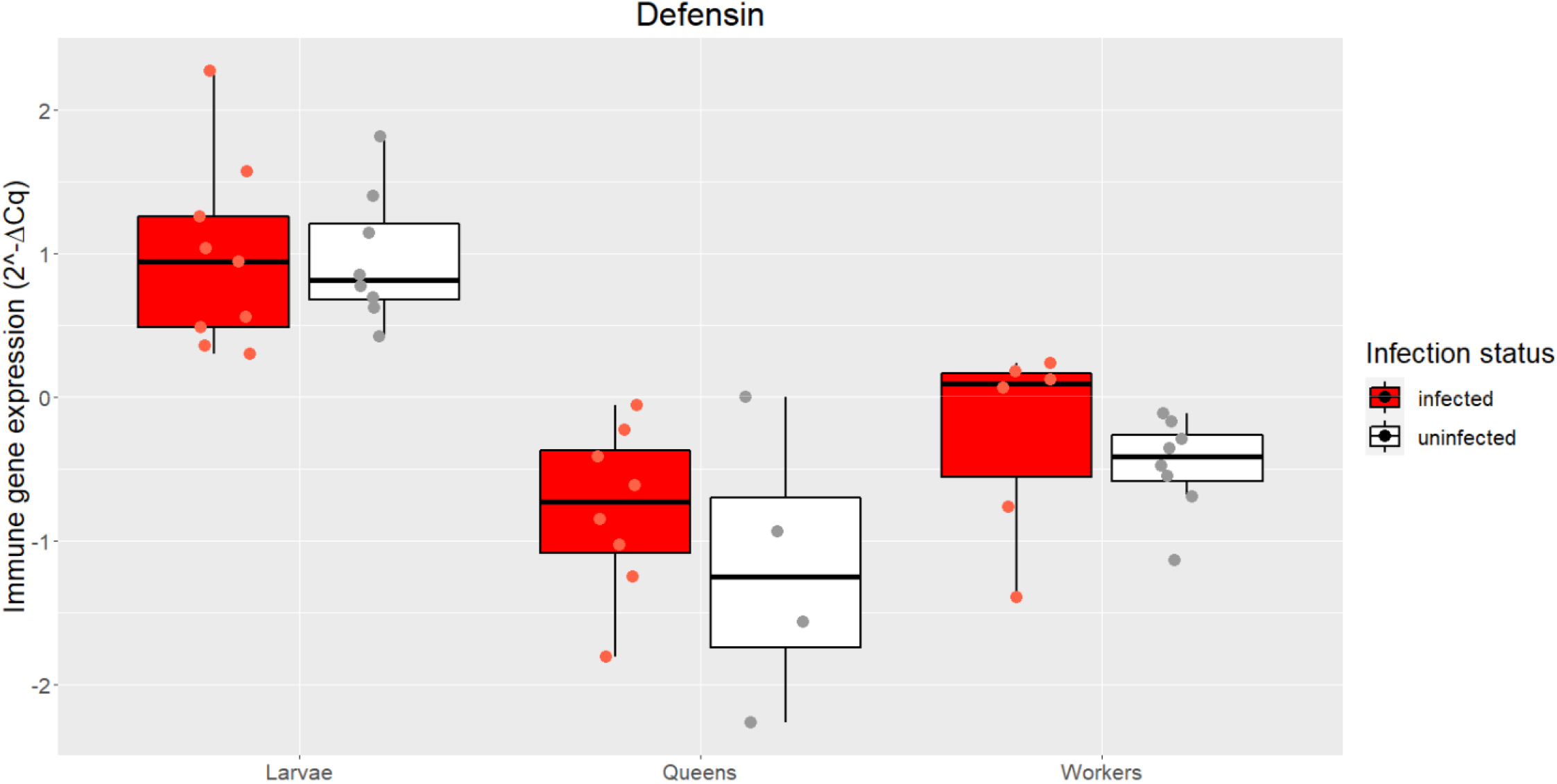
*Defensin 1* expression levels (2^-ΔCq^ values) in larvae, queens and workers from infected (red) and uninfected (white) colonies (transformed data according to the applied statistical analyses). Box-and-whisker plots are shown with median, 25% – 75% quartiles.

## Discussion

In social insects, investing energy and resources into mounting innate immune responses can divert resources from other essential functions, potentially affecting individual fitness and colony performance (Jacot et al. 2005; Ardia et al. 2012). We predicted that infection with a non-lethal parasite may lead to the upregulation of two key immune genes, *prophenoloxidase* and *defensin 1* and that infection effects on immune gene expression may vary between workers, queens and larvae due to their different exposure to infection risk and also considering differences in social roles.

Workers, being responsible for day-to-day tasks, such as foraging, nest maintenance, and brood care, face a high risk of pathogen exposure from the environment (Oster & Wilson 1978; Schmid-Hempel & Schmid-Hempel 1984). Consequently, their immune systems have evolved to effectively combat infections. Older foraging workers typically exhibit heightened cellular and humoral immune responses compared to young workers (Armitage & Boomsma 2010), which benefit nestmates, as workers are the main source of pathogen transfer into the colony (Armitage & Boomsma 2010; Bocher et al. 2007; Schmid et al. 2008; Schmid-Hempel 1998). Our study revealed that both *prophenoloxidase* and *defensin 1* level increased with pathogen loads in *M. scabrinodis* same-age workers. Our findings align with those reported by Konrad et al. (2012), who found heightened expression of both *prophenoloxidase* and *defensin 1* in *Lasius neglectus* ants exposed to the entomopathogenic fungi *Metarhizium anisopliae*. In a different study, Konrad et al. (2015) demonstrated that *Lasius neglectus* workers infected with *Metarhizium brunneum* and *Laboulbenia formicarum*, a close relative of *Rickia wasmannii*, showed upregulation of *prophenoloxidase* but no significant effect on *defensin 1*, even though wounding triggered *defensin 1* expression. *R. wasmannii* attaches to the surface of the cuticula of its ant host and perhaps initiates host immune defence mechanisms (e.g. upregulation of *prophenoloxidase and defensin* genes) in proportion to the number of thalli present. Surprisingly, this occurred independently of body size and fat content, suggesting that morphology and physiology of the host play only a minor, if any, role in innate immune responses of *M. scabrinodis* workers to non-lethal ectoparasitic fungi.

Queens are the key reproductives in the colony and prioritise different behavioural and physiological functions than non-reproductive workers (Hölldobler & Wilson 1990). Various studies indicate that immune activation in ant queens may affect egg-laying, lifespan, and even sperm viability (Gálvez & Chapuisat 2014; von Wyschetzki et al. 2016). Furthermore, it has been shown that *defensin 1* levels increase after mating, resulting in heightened pathogen resistance and an up-regulated immune system (Gálvez & Chapuisat 2014; Chérasse et al. 2019). Our results revealed a heightened level of phenoloxidase in highly infected queens, but no significant effect on *defensin 1*, i.e., queens exhibited a more pronounced immune response to a permanent, non-lethal infection. Studies in insects have also shown a link between immune function and body size with larger individuals displaying stronger immune responses (e.g., encapsulation immune response, Suwanchaichinda & Paskewitz 1998; Rantala et al. 2003). Our study revealed that queens with bigger heads exhibited elevated levels of *defensin 1* and queens with more fat content exhibited higher levels of *phenoloxidase*. This pattern might be attributed to the greater availability of physiological resources in larger queens, enabling them to allocate more towards immune defence.

While previous studies have uncovered the ability of adult social insects to develop immunity, limited research has delved into the immune function during the early life stages of insects. In our study, we explored whether the social context, specifically growing up in an infected colony, can trigger the activation of immune defence mechanisms in larvae. Unlike in adults, gene expression did not differ between larvae coming from infected or uninfected colonies. Other studies have found a significant up-regulation of *defensin 1* gene in *Apis mellifera* larvae infected with *Ascosphaera apis* chalkbrood fungus (Aronstein et al. 2010). However, in our case, the fungus *Rickia wasmannii* is found only in adult workers (Csata et al. 2014, Báthori et al. 2017) and is not known to infect ant larvae, though we expected that a stressful environment has a detrimental effect on larvae immune gene expression.

Our findings underscore the intricate nature of social insect immunity, suggesting that different caste members may employ tailored immune strategies to meet their specific physiological demands and priorities within the colony. Further exploration of these differences might enhance our comprehension of colony health and resilience in social insect societies.

## Author contributions

EC, KOB, BM, ES, and JH elaborated the study concept and design. Preparation of materials and data collection were performed by KOB, ES, BM, and EC. Data analyses were carried out by KOB, EC, and ES. Specific primers were designed by ES. Resources for experiments were provided by JH, EC and BM. The manuscript was written by KOB, EC, ES, JH, and BM. All authors read, contributed to and approved the final manuscript.

## Acknowledgements

We thank the Global Ant Genomics Alliance (GAGA) for providing us with the transcriptome of *Myrmica scabrinodis*. Alexander von Humboldt Foundation supported the study and EC’s work. KOB’s work was supported by the scholarship of Collegium Talentum, Hungary. We are grateful to Krisztina Puskás and Ágota Szabó for the field collection and to Georg Gebhard for his help with RNA extraction.

## Supplementary materials

**Table S1.** Detailed demographic data of the *Myrmica scabrinodis* colonies collected from the field for the study.

| Colony ID | Infection status of the colonies | Number of workers | Number of queens | Number of pupae | Number of larvae | Number of subnests |
| --- | --- | --- | --- | --- | --- | --- |
| FII | infected | 837 | 2 | 309 | 245 | 2 |
| FIII | infected | 823 | 11 | 367 | 359 | 8 |
| FIV | infected | 638 | 3 | 151 | 176 | 2 |
| FVI | infected | 1449 | 2 | 191 | 582 | 2 |
| NFI | uninfected | 736 | 6 | 78 | 288 | 2 |
| NFII | uninfected | 1141 | 6 | 206 | 407 | 6 |
| NFIII | uninfected | 944 | 1 | 81 | 444 | 1 |
| NFIV | uninfected | 1507 | 5 | 375 | 561 | 3 |
| NFV | uninfected | 1426 | 4 | 198 | 462 | 1 |

